# Neuronal mTORC1 inhibition promotes longevity without suppressing anabolic growth and reproduction in *C. elegans*

**DOI:** 10.1101/2021.08.12.456148

**Authors:** Hannah J. Smith, Anne Lanjuin, Arpit Sharma, Aditi Prabhakar, Emina Tabakovic, Rohan Sehgal, William B. Mair

## Abstract

One of the most robust and reproducible methods to prolong lifespan in a variety of organisms is inhibition of the mTORC1 (mechanistic target of rapamycin complex 1) pathway. mTORC1 is a metabolic sensor that promotes anabolic growth when nutrients are abundant. Inhibition of mTORC1 extends lifespan, but also frequently has other effects such as stunted growth, slowed development, reduced fertility, and disrupted metabolism. It has long been assumed that suppression of anabolism and resulting phenotypes such as impaired growth and reproduction may be causal to mTORC1 longevity, but this hypothesis has not been directly tested. RAGA-1 is an upstream activator of TORC1. Previous work from our lab using a *C. elegans* model of mTORC1 longevity, the long-lived *raga-1* null mutant, found that the presence of *raga-1* only in the neurons suppresses longevity of the null mutant. Here, we use the auxin-inducible degradation (AID) system to test whether neuronal mTORC1 inhibition is sufficient for longevity, and whether any changes in lifespan are also linked to stunted growth or fertility. We find that life-long AID of RAGA-1 either in all somatic tissue or only in the neurons of *C. elegans* is sufficient to extend lifespan. We also find that AID of RAGA-1 or LET-363/mTOR beginning at day 1 of adulthood extends lifespan to a similar extent. Unlike somatic degradation of RAGA-1, neuronal degradation of RAGA-1 doesn’t impair growth, slow development, or decrease the reproductive capacity of the worms. Lastly, while AID of LET-363/mTOR in all somatic cells shortens lifespan, neuronal AID of LET-363/mTOR slows aging. This work demonstrates that targeting mTORC1 specifically in the neurons uncouples longevity from growth and reproductive impairments, challenging previously held ideas about the mechanisms of mTORC1 longevity and elucidating the promise of tissue-specific aging therapeutics.

## INTRODUCTION

As humans grow older, they become susceptible to an increasing number of diseases that ravage quality of life. It is now generally accepted that the onset and rate of aging are not fixed. Decades of research have converged on nutrient metabolism as a potent regulator of aging^1^. One of the metabolic pathways that is most frequently targeted in aging studies is the mechanistic target of rapamycin complex 1 (mTORC1) signaling pathway^2,3^. mTORC1 is a multiprotein complex that senses energy and nutrient availability to promote growth when resources are abundant. mTORC1 incorporates information about the availability of insulin, growth factors, amino acids, energy, and cellular stress through upstream proteins like Rag GTPases and the tuberous sclerosis complex (TSC). When activated, mTORC1 promotes anabolic processes such as nucleotide and lipid synthesis while inhibiting catabolic processes such as autophagy to drive cellular growth and development^4^. Inhibition of the mTORC1 pathway, either genetically or pharmacologically, extends lifespan in yeast^5^, worms^6,7^, flies^8^, and mice. organisms ranging from yeast to mice^9–11^.

The success of mTORC1 inhibition in promoting the longevity of lab animals has made it a promising strategy for anti-aging interventions in humans, but it is important to consider that mTORC1 inhibition has many effects beyond lifespan extension. We have information on the effects of pharmacological mTORC1 inhibition in humans since certain mTORC1 inhibitors are approved for clinical use, mostly as immunosuppressants but also for the treatment of certain cancers. Administration of mTORC1 inhibitors to humans often results in decreased wound healing, reduced fertility, and, very commonly, metabolic disruptions like hyperglycemia and dyslipidemia^12^. While some of these pathologies may be resulting from off-target effects of the drugs, they also highlight the fact that the use of broad mTORC1 inhibitors as an anti-aging therapeutic is likely to disrupt other mTORC1 functions. In order to develop more targeted therapeutics that achieve longevity from mTORC1 inhibition while avoiding trade-offs, it would be beneficial to know whether there are specific tissues in which mTORC1 inhibition acts most strongly to counteract aging.

Genetic models of mTORC1 inhibition in worms, flies, and mice reveal another set of trade-offs reflective of the essential functions of the mTORC1 pathway. Genetic mutants with reduced mTORC1 signaling are long-lived but are often smaller, develop slower, and are less fertile^11,13,14^. Multiple theories posit that these disruptions in anabolic growth and development aren’t mere side effects but are actually causal to the lifespan extension that results from mTORC1 inhibition. These Y allocation theories draw on the idea that in the wild, animals have limited resources, so growth and reproduction occur at the expense of somatic maintenance. In times when resources are scarce, an organism allocates more energy to self-preservation, sacrificing growth and fertility, so that it can survive until conditions are more favorable to reproduce^15,16^. The observed phenotypes occurring after mTORC1 inhibition in the lab suggest that mTORC1 might be the central node that controls energy allocation and therefore extends lifespan by promoting the trade-off between somatic maintenance and fertility. However, this hypothesis has not been directly tested, and it is crucial to understand which, if any, of these trade-offs are causal to mTORC1 longevity in order to understand the mechanism of lifespan extension by mTORC1 and its therapeutic potential in humans.

Previous work from our lab has begun to address the tissue and trade-offs specific to mTORC1 longevity. The *C. elegans raga-1* null mutant is a genetic model of mTORC1 inhibition. *raga-1* encodes the ortholog of mammalian RAGA-1, an upstream regulator of mTORC1 that activates the complex when amino acids are present, and the *raga-1* null mutant is long-lived but also smaller, developmentally delayed, and less fertile. Rescuing expression of *raga-1* only in the neurons of the otherwise null animals suppresses the longevity, returning the animals to a wild type lifespan. In contrast, the neuronal expression of *raga-1* does not rescue the stunted body size or slowed development^14^. Altogether, these data suggest that mTORC1 regulates aging through the neurons and that this effect on aging may be independent of other phenotypes associated with whole-body mTORC1 inhibition. However, it was not known whether inhibition of mTORC1 only in the neurons would be sufficient to extend lifespan and whether this would impact peripheral phenotypes.

Our ability to determine the mechanisms driving mTORC1 longevity and discern which phenotypes can be uncoupled from mTORC1’s effect on aging have previously been limited by the methods most commonly used to inhibit mTORC1: genetic or pharmacological approaches that inhibit mTORC1 chronically or throughout the whole-body. Here, we use auxin-inducible degradation (AID) to achieve neuron-specific mTORC1 inhibition in *C. elegans*^17^. We find that lifelong degradation of RAGA-1 in the neurons or degradation of RAGA-1 or LET-363/mTOR from day 1 of adulthood onwards extends lifespan. We observe that lifelong degradation of RAGA-1 in all somatic tissues impairs growth, development, and reproduction as expected, but lifelong neuronal degradation doesn’t. Therefore, restricting mTORC1 inhibition to the neurons in *C. elegans* achieves lifespan extension without impairing growth or reproduction. This study reveals that mTORC1-mediated longevity does not require the suppression of growth or fertility and that mTORC1 acts in a specific tissue to regulate aging independently from other phenotypes.

## RESULTS

### Generation of strains for somatic and neuronal mTORC1 inhibition

We utilized the auxin-inducible degron (AID) system to target endogenous genes and achieve tissue-specific mTORC1 inhibition. The AID system, originally discovered in plants and adapted for protein degradation in *C. elegans*, offers multiple advantages over previous methods used to inhibit mTORC1: it is inducible, tissue-specific, and reversible and it allows for manipulation of the endogenous gene. In brief, TIR1 is a plant enzyme that interacts with endogenous *C. elegans* proteins to form an E3 ubiquitin ligase. In the presence of the plant hormone auxin, TIR1 binds degron-tagged proteins and ubiquitinates them, targeting them for degradation by the proteasome (Figure 1B).

**Figure 1.**
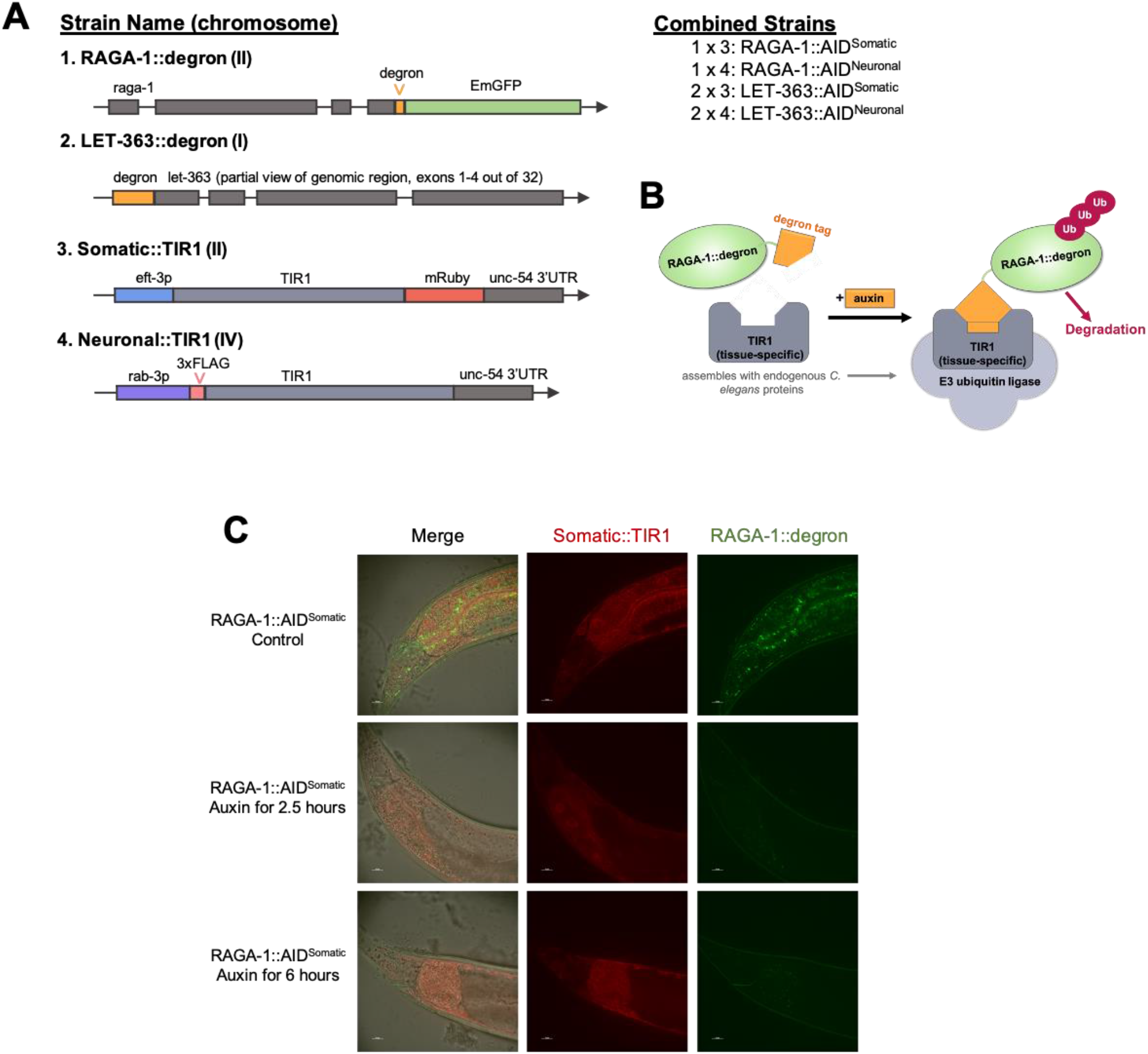
Schematic of tissue-specific auxin-inducible degradation (AID) of mTORC1 components. (A) Strains 1 and 2 were generated by CRISPR/Cas9 editing of the 3’ end of endogenous *raga-1* and the 5’ end of endogenous *let-363* with the 135 bp degron tag. Strain 3 is CA1200 obtained from CGC then backcrossed to our own N2 wild type strain (See Strains Table). The background *unc-119* mutation was eliminated. Strain 4 was generated by editing 3xFLAG-tagged TIR1 into the neuronal SKILODGE strain (Silva-García et al., *G3* 2019). (B) TIR1 assembles with endogenous *C. elegans* proteins to form an E3 ubiquitin ligase that recognizes and degrades degron-tagged proteins in the presence of the chemical auxin. (C) RAGA-1::AID^Somatic^ worms on control plates have TIR1 expression (red fluorescent) in all somatic tissues (not the germline) and RAGA-1::degron (green fluorescence) throughout all tissues, appearing as puncta. 2.5 hours after being moved onto 0.15 mM auxin plates, TIR1 is still visible but nearly all RAGA-1::degron is degraded in all tissues. 7 out of 21 imaged worms still had between 1 and 20 green puncta (see Figure 1C Supplemental). After 6 hours on 0.15 mM auxin, the GFP signal indicating RAGA-1::degron is very thoroughly diminished with only 6 out of 20 imaged animals containing 1 or 2 GFP puncta (see Figure 1C Supplemental). All images shown are of the tail region, see Figure 1C Supplemental for images of the midbody and germline. n = 1, at least 20 worms imaged per condition.

Using CRISPR/Cas9 gene editing, we tagged the 3’ end of endogenous *raga-1* with degron::EmGFP so that we could visualize expression and degradation of degron-tagged RAGA-1. We also tagged the 5’ end of endogenous *let-363* with the 135 bp degron sequence (Figure 1A0. The resulting alleles, *raga-1(wbm40)* referred to as RAGA-1::degron and *let-363(wbm46)* referred to as LET-363::degron, were verified by PCR and the strains were backcrossed 6 times to N2 to eliminate possible off-target mutations induced by Cas9.

The tissue in which RAGA-1::AID and LET-363::AID are degraded is determined by the expression of TIR1. Our lab previously generated a *C. elegans* gene expression toolkit called SKILODGE (single copy knock-in loci for defined gene expression) that allows for efficient CRISPR/Cas9-mediated knock-ins at the single copy level that are expressed in tissue-specific patterns^18^. We knocked TIR1 into the SKILODGE strain containing the *rab-3* promoter and the *rab-3* 3’ UTR to drive expression of TIR1 specifically in neurons, referred to as Neuronal::TIR1 (Figure 1A). For expression of TIR1 in all somatic tissue, we used a strain generated by another lab that drives TIR1 expression with the *eft-3* promoter, referred to as Somatic::TIR1^17^. We then crossed the RAGA-1::AID and LET-363::AID strains into the Neuronal::TIR1 and Somatic::TIR1 strains to achieve AID of mTORC1 components either in the whole body or in the neurons specifically (Figure 1A). Anecdotally, these strains had no obvious developmental or behavioral abnormalities relative to wild type animals.

### Efficient degradation is achieved at a low auxin concentration that minimizes off-target effects

Most AID experiments in *C. elegans* use concentrations of auxin around 1 mM. At this high of a concentration, auxin alone has effects on the animals such as promoting ER stress resistance^19^. To avoid this and any other potential side effects of high auxin concentrations, we performed an auxin concentration gradient experiment to find a low dose that was sufficient to induce degradation of an AID-tagged protein but without any effect on lifespan. We find that 0.15 mM auxin is sufficient to induce rapid and life-long degradation in a GFP::AID reporter strain (Figure 2 Supplemental A) without altering lifespan (Figure 2 Supplemental B). Alternatively, 1 mM auxin extends the lifespan of these animals (Figure 2 Supplemental B). Even at extremely low doses 0.01 mM and 0.025 mM, auxin is able to induce degradation of GFP::AID in day 1 animals, but this degradation is not maintained as the animals age (Figure 1 Supplemental A). Therefore, all subsequent experiments are performed with 0.15 mM auxin.

We imaged untreated and auxin-treated RAGA-1::AID^Somatic^ worms to test whether the RAGA-1::degron::EmGFP protein is thoroughly degraded with 0.15 mM auxin. At day 1 of adulthood, untreated animals have GFP signal in all tissues, mostly in the form of small puncta. These GFP-positive puncta are not present in wild type animals, therefore they most likely represent RAGA-1::degron::EmGFP protein. At day 1 of adulthood, RAGA-1::AID^Somatic^ animals were transferred to 0.15 mM auxin plates. After 2.5 hours, all GFP signal was diminished in a majority of the animals (Figure 1C), with 7 out of 20 animals still having between 1 and 20 puncta (Figure 1(C) Supplemental). After 6 hours on auxin, the GFP was thoroughly diminished in 14 out of 20 worms (Figure 1C), but 6 worms had 1 or 2 GFP puncta (Figure 1(C) Supplemental). These images demonstrate that 0.15 mM auxin efficiently and rapidly degrades degron-tagged RAGA-1.

### Lifespan extension by AID of RAGA-1 in soma and neurons

First, to confirm that RAGA-1 longevity using AID is possible, we tested the effect of whole-body and life-long AID of RAGA-1 using the RAGA-1::AID^Somatic^ strain with auxin administered from hatch. As expected, life-long, somatic degradation of RAGA-1 significantly extends lifespan (Figure 2A). Next, we used the RAGA-1::AID^Neuronal^ strain to test whether AID of RAGA-1 only in the neurons would be sufficient to promote longevity. Remarkably, neuronal RAGA-1 degradation is sufficient to significantly extend lifespan and to a similar extent as somatic degradation of RAGA-1 (Figure 2B). This suggests that neuronal mTORC1 is a potent regulator of aging in *C. elegans*.

**Figure 2.**
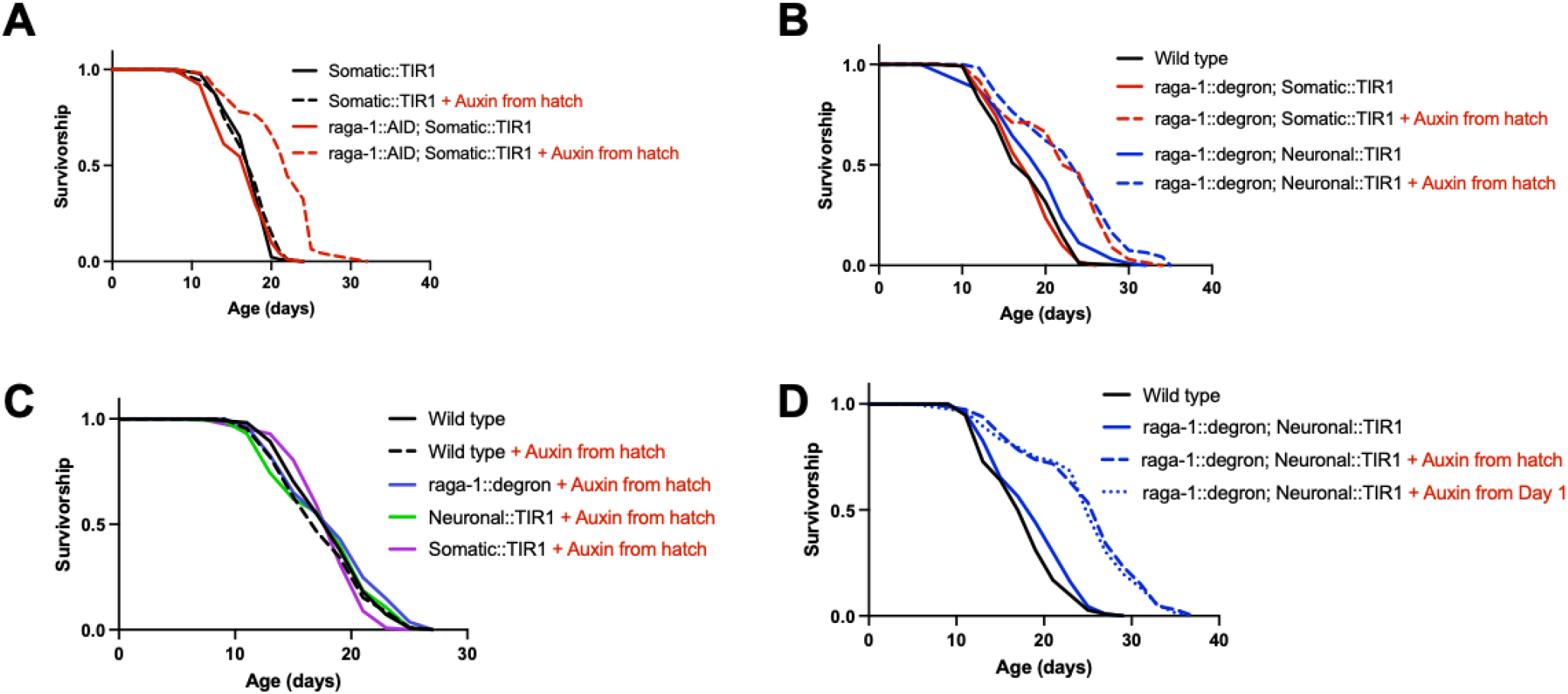
Auxin-inducible degradation of RAGA-1 in whole body or only in neurons extends lifespan. (A) Animals with life-long, whole body AID of RAGA-1 are significantly long-lived relative to the same animals on control plates (p<0.0001). n=4. (B) Worms with neuronal AID of RAGA-1 are significantly long-lived relative to the same animals on control (p<0.0001). Again, somatic AID of RAGA-1 extends lifespan (p<0.0001). There is no difference between worms with neuronal or somatic AID of RAGA-1 (p=0.2583), demonstrating that neuronal RAGA-1 degradation extends lifespan to a similar extent as whole-body degradation. (C) Life-long exposure to auxin doesn’t alter the lifespan of wild type worms (p=0.4515), worms expressing raga-1::degron only (p=0.2790), worms expressing Neuronal::TIR1 only (p=0.9632), or worms expressing Somatic::TIR1 only (p=0.2494). n=2 (D) Adult-onset AID of RAGA-1 in the neurons, beginning at day 1 of adulthood, is sufficient to extend lifespan (p<0.0001) and extends lifespan to the same extent as life-long AID of RAGA-1 in neurons (p=0.7100 between adult-onset and auxin from hatch conditions, ns). n=2.

Given that the AID system has been found to regulate the mTORC1 pathway in plants and fungi^20,21^, we analyzed the lifespan of numerous control strains to test whether the observed lifespan extension was truly due to RAGA-1 degradation rather than any off-target effects of auxin, the AID tag, or TIR1. We treated wild type animals, the RAGA-1::degron strain, and both TIR1 strains with lifelong auxin and they all had the same lifespan as untreated wild type animals (Figure 2C). Therefore, it is unlikely that auxin itself, the presence of TIR1 alone, or disruption of RAGA-1 function by the AID tag are contributing to the observed longevity phenotype. These data also suggest that there are no effects of auxin in combination with TIR1 alone or RAGA-1::degron alone. Additionally, when untreated with auxin, the RAGA-1::AID^Somatic^ and RAGA-1::AID^Neuronal^ strains have the same lifespan as wild type animals (Figure 2B). Thus, there is no evidence that the observed longevity is due to any auxin-independent interaction between TIR1 and RAGA-1::degron. Together, our data demonstrate that we have a robust model of longevity in *C. elegans* that is driven by neuron-specific AID of RAGA-1.

### Targeting RAGA-1 in the neurons uncouples longevity from other mTORC1 phenotypes

In addition to lifespan extension, whole-body mTORC1 inhibition has organismal consequences such as impaired growth, slowed development, and decreased reproductive capacity^7,13,14^. It has long been assumed that these phenotypes are causal to and therefore inseparable form mTORC1 longevity, but this hypothesis has gone untested. Our model of tissue-specific mTORC1 longevity provides us with an opportunity to test whether mTORC1 regulates any or all of these phenotypes through the neurons and whether these phenotypes are required for mTORC1-mediated lifespan extension. We measured body size, development time, and brood size in long-lived animals that had either somatic or neuronal RAGA-1 degradation from hatch and compared them to wild type animals.

Development time was measured as the time from when an egg was laid to when that animal grew up and laid its first egg. Body size was measured as the length of the worms on day 1 of adulthood. Finally, reproductive capacity was measured as the total number of eggs a worm laid over its lifetime. We find that animals with somatic degradation of RAGA-1 from hatch take significantly longer to develop (mean = 74.57 hours), but animals with neuronal RAGA-1 degradation (mean = 66.23 hours) develop in the same amount of time as wild type animals (mean = 65.98 hours) (Figure 3A, Figure 3(A&C) Supplemental). Similarly, animals with somatic RAGA-1 degradation lay significantly fewer eggs (mean = 183 eggs), but animals with neuronal RAGA-1 degradation (mean = 286 eggs) have a similar reproductive capacity to wild type animals (mean = 306 eggs) (Figure 3B). Finally, although animals with somatic AID of RAGA-1 from hatch are significantly shorter than wild type animals, animals with neuronal AID of RAGA-1 have a normal body size (Figure 3C, Figure 3(C) Supplemental, Figure 3(A&C) Supplemental).

**Figure 3.**
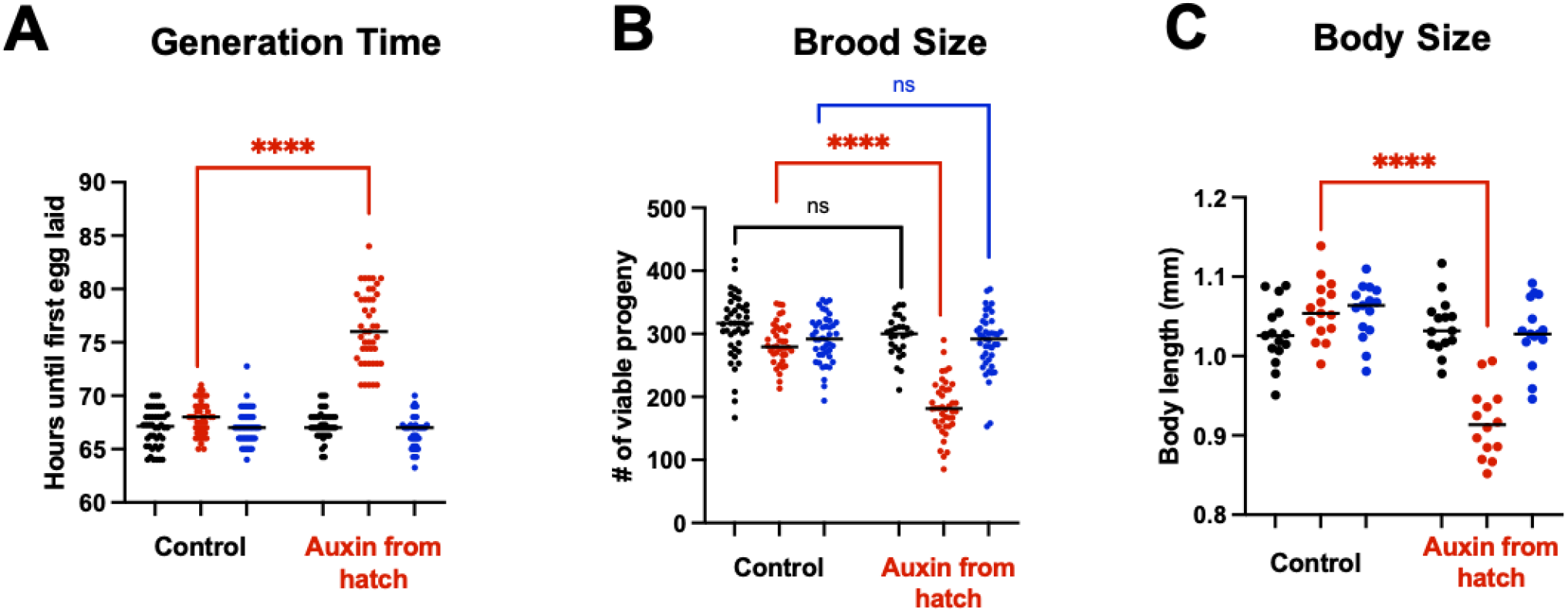
Neuronal RAGA-1 loss doesn’t impair generation time, brood size, or body size. (A) AID of RAGA-1 in all somatic tissue (raga-1::degron; Somatic::TIR1 + Auxin) results in significantly delayed generation time (p<0.0001). Alternatively, AID of RAGA-1 in neurons has no effect on generation time. There were no significant differences between the control-treated strains. n=3 independent replicates with at least 40 animals per group. (B) AID of RAGA-1 in the soma results in animals that have fewer viable offspring (p<0.0001) but animals with AID of RAGA-1 in the neurons have a wild type brood size. The only observed difference amongst the control-treated strains is that raga-1::degron, Somatic::TIR1 animals have a slightly smaller brood size than wild type animals (p = 0.0184). n=3 independent replicates with at least 40 animals except for wild type on auxin which only has 2 replicates with 28 animals total. (C) AID of RAGA-1 in somatic tissues results in significantly smaller body size at Day 1 of adulthood (p<0.0001). No differences were observed between control-treated strains. n=1 replicate with at least 14 animals per group. Additional replicate shown in figure supplement.

These findings have multiple implications for mTORC1 longevity. First and foremost, the fact that animals with neuronal AID of RAGA-1 are long-lived but have normal growth, development, and reproduction demonstrates that these suppression of these anabolic phenotypes can indeed be uncoupled from longevity. Moreover, these data shed new light on tissue-specific functions of mTORC1. While RAGA-1 is able to regulate aging from the neurons of *C. elegans*, it must regulate whole body size, development rate, and reproductive capacity through non-neuronal tissues.

### Lifespan extension by AID of LET-363/mTOR in neurons

To test whether this neuronal control of longevity is specific to RAGA-1 or applies more broadly to the mTORC1 pathway, we tested the effects of neuronal and somatic AID of LET-363/mTOR. Previously, there was not a good method to test the lifespan effects of LET-363 manipulation for two reasons. First, deletion of *let-363* is lethal, making tissue-specific rescue experiments implausible. Second, targeting *let-363* with RNA interference (RNAi) leads to confounding results because the RNAi also knocks down a mitochondrial gene, *mrpl-47*, that exists in an operon with *let-363* and had been found to affect lifespan^22,23^. Therefore, the tissue-specificity and inducibility of the AID system is particularly advantageous here and allows us to assess the relationship between LET-363 and aging with unprecedented specificity.

AID of LET-363 in all somatic tissues from hatch results in arrest at the L1 larval stage (Figure 4A). To avoid larval arrest, we tested the effect on lifespan of adult-onset LET-363 degradation in either the soma or only in the neurons. We find that whole-body degradation of LET-363 from day 1 onwards shortens the lifespan of the animals (Figure 4B). On the other hand, degradation of LET-363 only in the neurons extends lifespan (Figure 4B). These data demonstrate that LET-363 inhibition is deleterious in some tissues but advantageous in others, highlighting the importance of identifying the key tissue(s) in which a pathway acts to regulate aging rather than targeting it broadly throughout the entire body.

**Figure 4.**
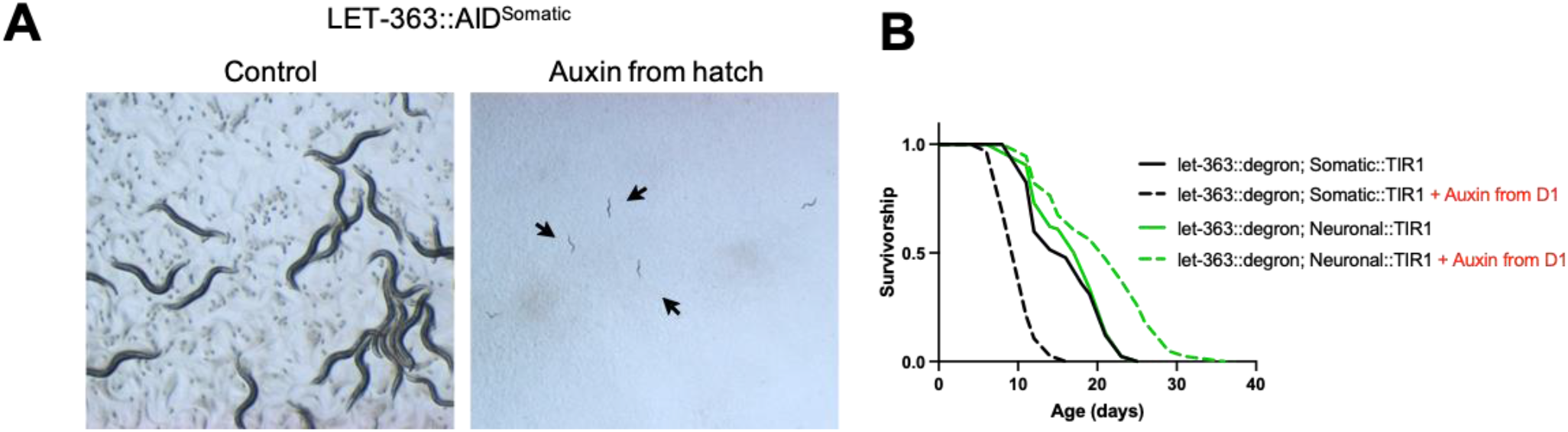
Somatic AID of LET-363 causes larval arrest or shorter lifespan, but adult-onset neuronal AID of LET-363 extends lifespan. (A) 3 days after eggs are laid, LET-363::AID^Somatic^ animals on control (left) have grown into gravid Day 1 adults. Animals grown on 0.15 mM auxin since hatch (right) are arrested at the L1 larval stage. Both imaged at 1.6x. (B) Lifespan of LET-363::AID^Somatic^ animals that were hatched on control plates and then moved to either control plates or plates containing 0.15 mM auxin on day 1 of adulthood. Degradation of LET-363::degron in all somatic tissues beginning at day 1 of adulthood results in decreased lifespan (p<0.0001). Alternatively, adult-onset degradation of LET-363 only in the neurons significantly extends lifespan (p<0.0001). n=1 for neuronal versus somatic comparison. n=2 for lifespan extension by neuronal AID of LET-363. n=2 for lifespan shortening by whole-body AID of LET-363, but one replicate used a lower concentration (0.025 mM) auxin.

## DISCUSSION

Decades of research into the biological pathways that regulate the rate of aging have converged on nutrient metabolism pathways such as mTORC1. Animal studies have revealed that inhibition of the mTORC1 pathway holds much promise as a strategy to promote healthy aging. However, translation of mTORC1 therapeutics into humans is hindered by the fact that whole-body mTORC1 inhibition also induces undesirable effects such as reduced fertility, stunted growth, metabolic disruption and immunosuppression. One potential strategy is to identify the key tissue(s) in which mTORC1 acts to regulate aging and then design targeted therapies that promote healthy aging without unwanted side effects.

Here, we find that neuron-specific mTORC1 inhibition is sufficient for longevity in *C. elegans* using AID, a robust system with multiple advantages over previous methods used to study the tissue-specific effects of mTORC1 on aging. In *C. elegans*, intestine-specific RNAi of *ragc-1* was shown to extend lifespan^7^. In flies, lifespan was extended by manipulation of mTORC1 pathway components using GAL4-UAS drivers primarily expressed in the fat body^8^. However, further research has shown that both tissue-specific RNAi^24^ and GAL4-UAS drivers have off-target effects in other tissues^25^. In addition to being more tissue-specific than RNAi, AID has the added benefit of targeting the endogenous protein directly rather than silencing a gene at the mRNA level. Therefore, late-onset AID is able to eliminate any of the target protein that is present while late-onset RNAi is unable to eliminate protein but is only able to prevent more from being translated.

Researchers have taken advantage of the AID system to uncover new knowledge about another major longevity pathway in *C. elegans*: the insulin/insulin-like growth factor 1 (IGF-1) signaling (IIS) pathway. The researchers were able to begin AID of DAF-2, ortholog of mammalian insulin and IGF-1 receptor, at day 25 adulthood, a time when RNAi of DAF-2 is not effective. They found that AID of DAF-2 late in life, or whole-life AID only in the intestine or neurons, extended lifespan in *C. elegans*^26^. Future studies using the AID system to identify the key tissues in which nutrient metabolism pathways act to regulate aging will shed new light on the specific mechanisms driving the resulting longevity.

Our finding that neuronal mTORC1 inhibition extends lifespan without impairing growth, development, or fertility challenge many of the current theories about the mechanisms driving mTORC1 longevity. It has long been assumed that trade-offs are causal to mTORC1 longevity and that mTORC1 inhibition extends lifespan by shunting resources away from growth or reproduction and back into somatic maintenance. Indeed, there is a strong association between small body size and longevity that isn’t unique to mTORC1 inhibition, but is also seen between animals of the same species and in long-lived animals with mutations in the insulin and growth hormone signaling pathways^27^. However, we find that animals with neuron-specific mTORC1 inhibition are long-lived but are the same size and have the same reproductive capacity as wild-type animals. This suggests that mTORC1 regulates aging through a different tissue, in this case the neurons, than reproduction or growth at the whole-body level. Therefore, neither a reduction in body size or in reproduction is required for mTORC1 longevity. Lastly, that somatic degradation of LET-363 shortens lifespan while neuronal degradation increases it highlights the fact that the same enzyme can have both pro and anti-aging phenotypes in a tissue specific manner. This concept has far-reaching consequences for anti-aging therapeutics that are taken orally and affect a target across multiple tissues.

Overall, this research demonstrates the importance of determining whether an intervention is acting throughout the whole-body or only in specific tissues to promote healthy aging. Tissue-specific interventions may allow for more translational anti-aging strategies with fewer negative side effects. Additionally, the development of models of tissue-specific longevity, such as the neuron-specific model of mTORC1 longevity presented here, present a unique opportunity to identify mechanisms of the targeted pathway that specifically mediate longevity.

## METHODS

### *C. elegans* strains and husbandry

Worms were maintained at 20°C on nematode growth media (NGM) plates seeded with *E. coli* strain OP50-1(CGC) using standard techniques^28^. *E. coli* were grown in LB in a shaker at 37°C overnight, then 100 μL of the liquid culture was seeded onto the plates and left for 2 days to grow at room temperature.

| Strain | Genotype | Notes* | Source |
| --- | --- | --- | --- |
| N2 Bristol | Wild type |  | CGC |
| WBM1204 | *raga-1(wbm40) [raga-1::degron::EmGFP] II* | **RAGA-1::degron** | Mair Lab |
| WBM1481 | *let-363(wbm46) [degron::let-363] I* | **LET-363::degron** | Mair Lab |
| WBM1223 | *wbmls83 [rab-3p::3xflag::TIR1::rab-3 3'UTR \*wbmls66] IV* | **Neuronal::TIR1** | Mair Lab |
| CA1200 | *ieSi57 [eft-3p::TIR1::mRuby::unc-54 3'UTR + Cbr-unc-119(+)] II; unc-119(ed3) III* | **Somatic::TIR1** | CGC |
| WBM1395 | *ieSi57 [eft-3p::TIR1::mRuby::unc-54 3'UTR + Cbr-unc-119(+)] II* | CA1200 backcrossed 6 times to N2 |  |
| WBM1138 | *raga-1(wbm40) [raga-1::AID::EmGFP], ieSi57 [eft-3p::TIR1::mRuby::unc-54 3'UTR + Cbr-unc-119(+)] II* | **RAGA-1::AID<sup>Somatic</sup>**, 4x outcrossed *raga-1(wbm40)* crossed to CA1200 | Mair Lab |
| WBM1438 | *raga-1(wbm40) [raga-1::AID::EmGFP], ieSi57 [eft-3p::TIR1::mRuby::unc-54 3'UTR + Cbr-unc-119(+)] II* | **RAGA-1::AID<sup>Somatic</sup>**, WBM1204 x WBM1395, | Mair Lab |
| WBM1250 | *raga-1(wbm40) [raga-1::degron::EmGFP] II; wbmls83 [rab-3p::3xflag::TIR1::rab-3 3'UTR \*wbmls66] IV* | **RAGA-1::AID<sup>Neuronal</sup>** | Mair Lab |
| CA1202 | *ieSi57 [eft-3p::TIR1::mRuby::unc-54 3'UTR + Cbr-unc-119(+)] II; ieSi58 [eft-3p::degron::GFP::unc-54 3'UTR + Cbr-unc-119(+)] IV* |  | CGC |
| WBM1137 | *let-363(wbm46) [AID::let-363] I; ieSi57 [eft-3p::TIR1::mRuby::unc-54 3'UTR + Cbr-unc-119(+)] II* | **LET-363::AID<sup>Somatic</sup>**, *let-363(wbm46)* crossed to CA1200, not outcrossed | Mair Lab |
| WBM1249 | *let-363(wbm46) [AID::let-363] I; wbmls83 [rab-3p::3xflag::TIR1::rab-3 3'UTR \*wbmls66] IV* | **LET-363::AID<sup>Neuronal</sup>**, *let-363(wbm46)* crossed to WBM1250 | Mair Lab |

\*All Mair Lab strains are 6x outcrossed unless otherwise stated

### Microinjection and CRISPR/Cas9 gene editing

CRISPR-mediated gene editing was performed according to^29^. For the Neuronal::TIR1 strain, the TIR1 CRISPR mix was injected into the Neuronal SKILODGE strain as described in PMID: 31064766. Briefly, homology repair templates were amplified by PCR, using primers that introduced a minimum stretch of 35 bp homology at both ends. Single-stranded oligo donors (ssODN) were also used as repair templates. CRISPR injection mix reagents were added in the following order: 0.375 μl Hepes pH 7.4 (200 mM), 0.25 μl KCl (1 M), 2.5 μl tracrRNA (4 μg/μl), 0.6 μl *dpy-10* crRNA (2.6 μg/μl), 0.25 μl *dpy-10* ssODN (500 ng/μl), and PCR or ssODN repair template(s) up to 500 ng/μl final in the mix. If the edit was on Chromosome II, *dpy-5* was used as a coinjection marker. Water was added to reach a final volume of 8 μl. 2 μl purified Cas9 (12 μg/μl) added at the end, mixed by pipetting, spun for 2 min at 13000 rpm and incubated at 37 °C for 10 min. Mixes were microinjected into the germline of day 1 adult hermaphrodite worms using standard methods^30^. Alleles were verified by PCR and strains and most strains were backcrossed to N2 wild type worms 6 times before use to eliminate off-target mutations. Strain outcross information can be found in the table above.

### Pouring auxin and control plates

A 400 mM stock of indole-acetic acid (auxin) was prepared in ethanol, filter-sterilized, and stored at 4°C in a foil-covered tube as described in^17^. A batch of NGM (with or without Carbenicillin) was prepared and then divided. In one half, 0.15 mM auxin was added to the NGM by diluting the 400 mM auxin stock in either M9 or sterile Milli-Q water. Control plates were poured by diluting an equivalent amount of 100% filter-sterilized ethanol in M9 or water and then adding it to the other half of NGM. Auxin and control plates were covered in a black tarp or foil to dry overnight before being moved to 4°C. Plates were taken out and covered one night prior to seeding with a culture of HT115 bacteria.

For some experiments, instead of auxin being mixed into the media before the plates were poured, auxin or control solutions were pipetted on top of the plate. In this case, 100 uL of 15 mM auxin was pipetted onto seeded plates the evening before they were to be used for the experiment. Given that the plates contain 10 mL of media, diffusion of the auxin would result in a final concentration of 0.15 mM. Control plates were prepared by pipetting 100 uL of a solution that has an equivalent amount of ethanol.

### Generation time assay

Day 1 gravid adults were bleached and eggs were pipetted onto 6 cm NGM plates containing 100 μg/mL Carbenicillin seeded with *E. coli* strain HT115 bacteria. Animals were synchronized by performing an egg lay in the evening 3 days prior to the experiment, except the egg lay of RAGA-1::AID^Somatic^ animals onto auxin was performed that morning due to their developmental delay. For the egg lays, 10-20 animals were placed on a control or auxin plate and allowed to lay eggs for 45 minutes and were then removed. Early in the morning 3 days later, 15 L4 animals for each condition were picked onto individual 3.5 cm NG Carb plates that were 1-day seeded with 50 μL of HT115 bacteria. Animals were scored every hour until the first egg was laid.

### Brood size measurement

Animals were bleached onto NG Carb plates and synchronized by egg lay as mentioned above in “Generation time assay”. 15 animals from each condition were singled out onto individual 3.5 cm NG Carb plates seeded with HT115 bacteria at the L4 stage. For the next two days, animals were transferred both in the morning and in the evening to freshly seeded plates. For the following two days, animals were transferred only once. The plates containing the eggs were maintained in the incubator at 20°C for 3 days until the progeny had grown into day 1 adults. The number of progeny on each plate were counted and the counts from each day were summed for each individual parent.

### Body size measurement

Worms were anesthetized on an auxin or control plate without bacteria using 1 mg/mL tetramisole/M9. Once still, worms were imaged on a Zeiss Discovery V8 microscope with Axiocam camera. All animals were imaged in brightfield at 8x with a constant exposure. At least 12 worms were imaged per condition per replicate. Body length was analyzed in ImageJ by drawing a line end-to-end down the middle of the worm and measuring the length of the line.

### Lifespans

Lifespans were conducted at 20°C on 6 cm NGM plates with either 10 or 20 worms per plate and starting with 100-120 worms per condition. Lifespans were performed as described in^31^.

### Statistical analysis

Data was graphed and analyzed in GraphPad Prism 9. For lifespan experiments, survival curves were analyzed using the Log-rank test. For generation time, brood size, and body size, outliers were excluded using the ROUT method. A one-way ANOVA followed by a multiple comparisons test was performed on the cleaned data set to determine which groups were statistically significant different relative to the N2 control group.

---

**Figure 1(C) Supplemental.**
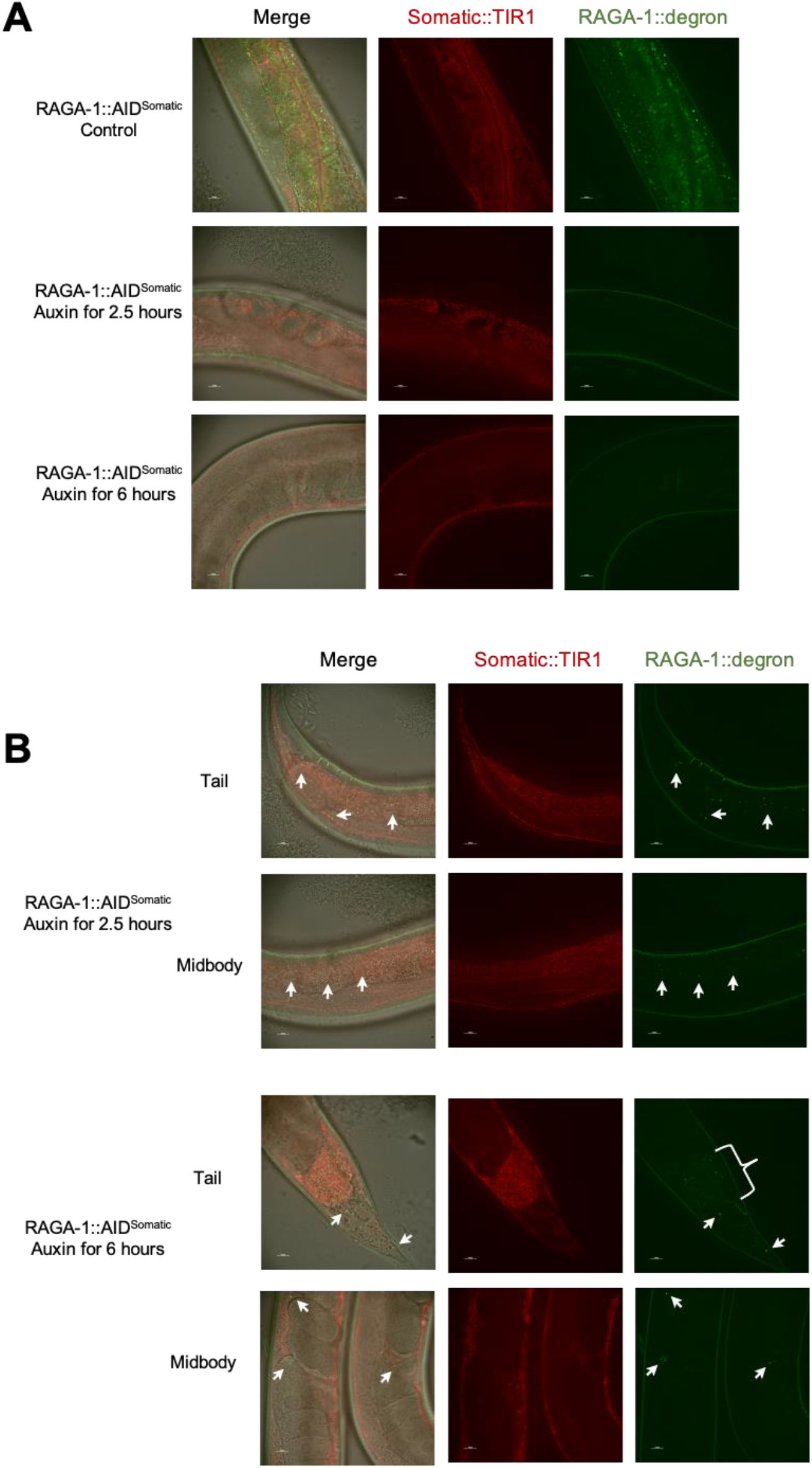
Images of midbody and representative images with partial AID of RAGA-1::degron. (A) Representative images of the midbody region. (B) Representative images of the worms with partial degradation of RAGA-1::degron at the corresponding time points. White bracket indicates intestinal autofluorescence. Picture in last row shows two worms side-by-side.

**Figure 2 Supplemental.**
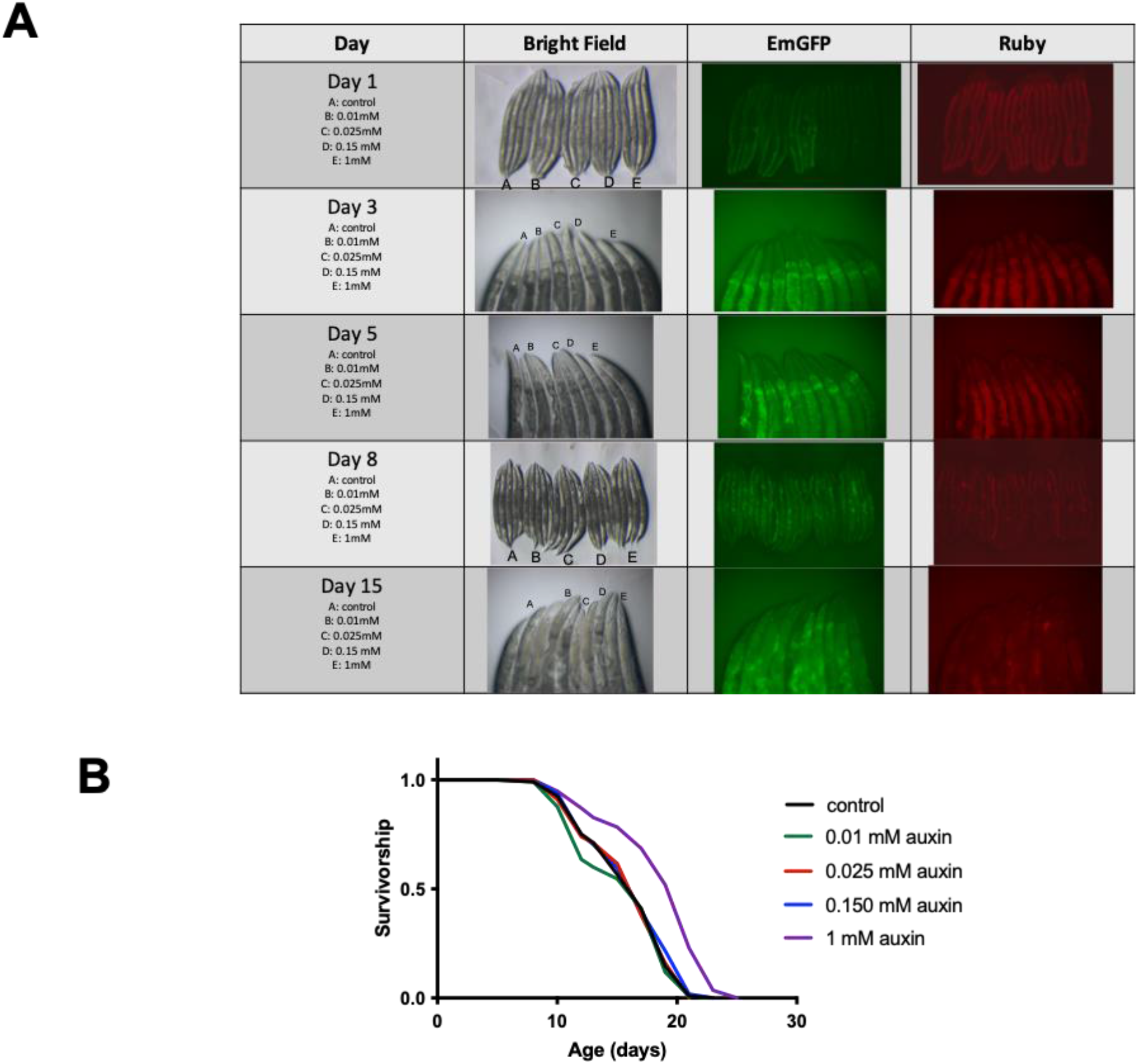
0.15 mM auxin is sufficient to degrade degron::GFP without affecting lifespan. Animals expressing eft-3p::TIR1::mRuby and eft-3p::degron::GFP were exposed to various concentrations of auxin and imaging and lifespan analyses were performed. (A) TIR1::mRuby is stably expressed across lifespan. 0.15 mM and 1 mM are the only concentrations of auxin that degraded degron::GFP even at Day 15 of adulthood. Images at later days were focused on the heads of *C. elegans* due to the intensity of green autofluorescence in the intestine. (B) *C. elegans* on 1 mM of auxin live significantly longer than animals of the same genotype on control plates (p<0.0001), but the lifespan of animals administered 0.15 mM auxin is unaffected. n=1.

**Figure 3(A&C) Supplemental.**
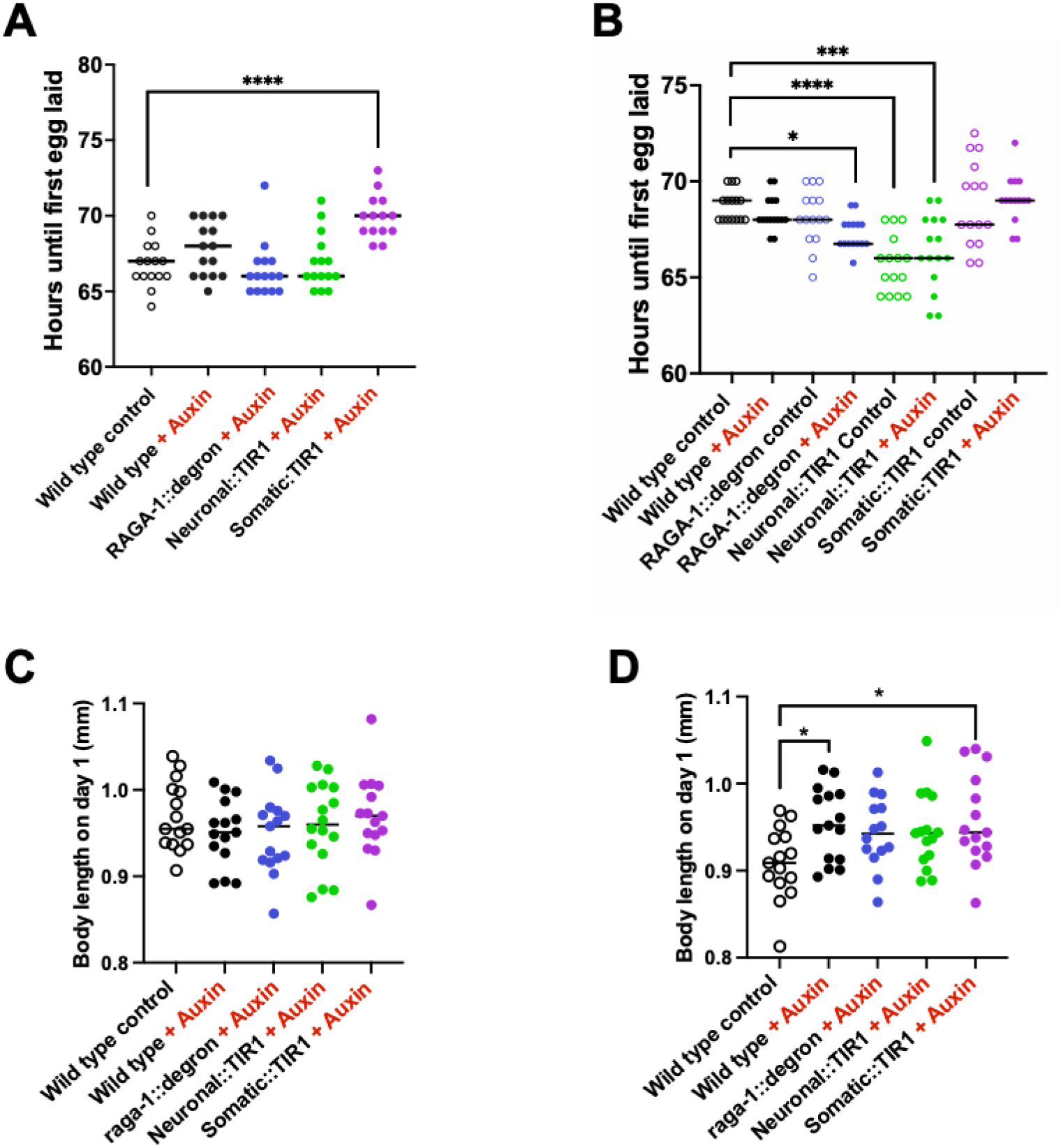
Control strains for generation time and body size. For all auxin conditions, 0.15 mM auxin was administered from hatch. (A & B) Generation time of control strains, RAGA-1::degron alone or Somatic or Neuronal::TIR1 alone. Some strains were found to have statistically significant generation times than the wild type control condition, but none of the significant differences held across the both replicates. n = 2, at least 15 animals per condition. (C & D) The body size of control strains was analyzed. In one of the replicates, wild type animals on auxin (p = 0.0231, median = 0.909) and Somatic::TIR1 animals on auxin (p = 0.0135, median = 0.944) were found to be slightly larger than wild type control animals (median = 0.909). n =2, at least 14 animals per condition.

**Figure 3(C) Supplemental.**
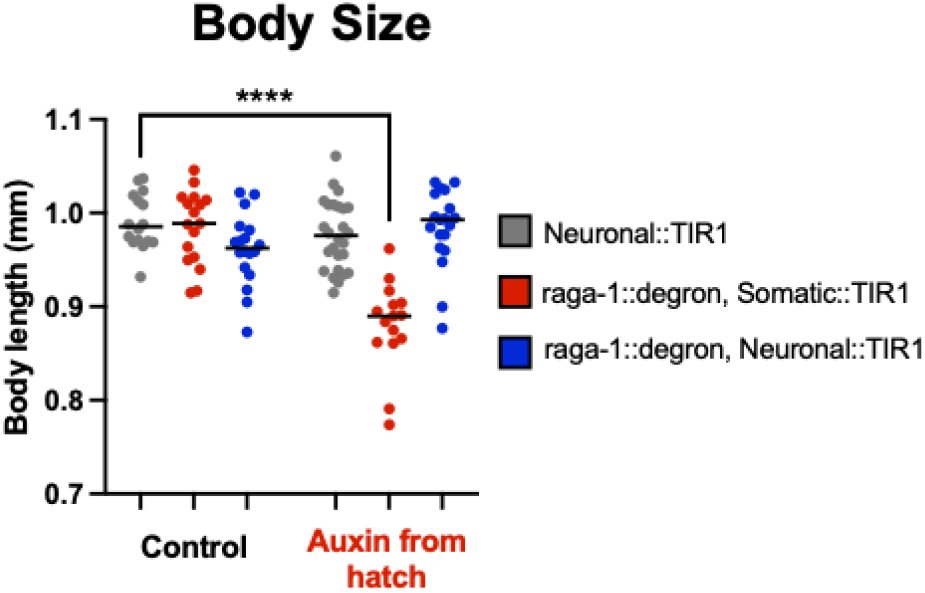
Repeat of body size with Somatic::TIR1 not outcrossed. Body size experiment repeated as in Figure 5C but with Somatic::TIR1 not backcrossed to our lab’s own N2 wild type worms (WBM1138 here as opposed to WBM1438 used in Figure 5C, see Strains Table for more information).

